# Multi-Level Contrastive Learning for Protein-Ligand Binding Residue Prediction

**DOI:** 10.1101/2023.12.06.570503

**Authors:** Jiashuo Zhang, Ruheng Wang, Leyi Wei

**Author notes:** Corresponding authors: L.W.

## Abstract

Protein-ligand interactions play a crucial role in various biological functions, with their accurate prediction being pivotal for drug discovery and design processes. Traditional methods for predicting protein-ligand interactions are limited. Some can only predict interactions with a specific molecule, restricting their applicability, while others aim for multiple types but fail to effectively utilize information across different interactions, leading to increased complexity and inefficiency. This study presents a novel deep learning model named MucLiPred and a dual contrastive learning mechanism aimed at improving the prediction of multiple ligand-protein interactions and the identification of potential ligand-binding residues. We proposed two novel contrastive learning paradigms at residue and type levels, training the discriminative representation of samples. The residue-level contrastive learning hones in on distinguishing binding from non-binding residues with precision, shedding light on nuanced local interactions. In contrast, the type-level contrastive learning delves into the overarching context of ligand types, ensuring that representations of identical ligand types gravitate closer in the representational space and bolstering the model’s proficiency in discerning interaction motifs, enhancing the model’s ability to recognize global interaction patterns. This approach culminates in nuanced multi-ligand predictions, unraveling relationships between various ligand types, and fortifying the potential for precise protein-ligand interaction predictions. Empirical findings underscore MucLiPred’s dominance over existing models, highlighting its robustness and unparalleled prediction accuracy. The integration of dual contrastive learning techniques amplifies its capability to detect potential ligand-binding residues with precision. By optimizing the model’s structure, we discovered that separating representation and classification tasks, leads to improved performance. Consequently, MucLiPred stands out as a groundbreaking tool in protein-ligand interaction prediction, laying the groundwork for future endeavors in this complex arena.

## 1 Introduction

Protein binding plays a crucial role in the functioning of living organisms as it encompasses molecular interactions involving DNA–protein, RNA–protein, and peptide– protein. These interplays contribute to a multitude of biological functions within the cellular structures of these organisms. Numerous biological processes, including gene expression regulation, signal transduction, and post-transcriptional modification and regulation, are fundamentally influenced by interactions between proteins and nucleic acids (DNA or RNA). Consequently, precise identification of DNA– and RNA-binding residues becomes indispensable for both the development of innovative therapeutics and the examination of mechanisms underlying protein-nucleic acid interactions. Interactions between DNA and proteins serve as a cornerstone for an array of biological actions, spanning transcription, regulation of gene expression, and splicing ^1, 2^. Additionally, protein-RNA interactions are integral to a multitude of biological operations, including the regulation of gene expression, viral assembly and replication, post-transcriptional modifications, and the synthesis of proteins ^3–5^. Moreover, protein-peptide interactions, being one of the most critical forms of protein associations, are instrumental in various cellular operations, including, but not limited to, DNA repair and replication, gene expression, and metabolic processes ^6, 7^. For a comprehensive understanding of the functions and mechanisms in any biological operation, it becomes essential to gain insight into the interaction residues between proteins and various molecules ^8^. Therefore, determining the residues interacting with various molecules can aid in biotechnological manipulation. Advancements in experimental techniques, including X-ray crystallography and nuclear magnetic resonance (NMR), have facilitated the discovery and documentation of multiple protein-ligand interacting residue structures, which have been reported and archived in the Protein Data Bank (PDB) ^9^. Unfortunately, due to the limitations of structure determination techniques, only a fraction of protein structures involved in interactions with other molecules have been deposited in the PDB. Additionally, these experimental methods are highly costly and time-consuming. In contrast, computational approaches that rely on sequence information offer a highly viable and cost-efficient alternative for the potential identification of binding residues and other biochemical problems ^10–13^. Hence, the development of computational methodologies for predicting protein-ligand binding residues is of utmost urgency ^14^.

Over recent years, an extensive assortment of computational algorithms has emerged for the forecasting and discernment of proteins and residues interacting with various molecules. In general, these methods can be classified into two main categories, single-type prediction and multi-type prediction methods. In the case of single-type prediction methods, the model is capable of predicting the interacting residues between proteins and a single type of molecule. For DNA-binding residues, various methods have been developed. Amirkhani et al. created a predictor called funDNApred ^15^ for DNA-binding residues in protein sequences using the Fuzzy Cognitive Map (FCM) model, leveraging information about solvent accessibility, evolutionary conservation, and amino acid propensities. Zhu et al. advanced the Ensembled Hyperplane-Distance-Based Support Vector Machines (E-HDSVM) for protein-DNA binding site prediction, implemented in DNAPred ^16^. Additionally, Patiyal et al. proposed a hybrid model called DBPred ^17^ using both traditional machine learning and deep learning, combining various features to forecast DNA-protein interactions. For RNA-binding residues, there are many methods have been proposed. Kumar et al. ^18^ introduced a model grounded on sequence data, utilizing machine learning strategies and evolutionary information. Walia et al. developed RNABindRPlus ^19^, a computational model that combines sequence homology-based methods with an optimized Support Vector Machine classifier, enhancing the reliability of predicting RNA-binding residues in proteins. Furthermore, Shen et al. developed RPiRLS ^20^, a machine-learning method that integrates a sequence-based derived kernel with regularized least squares, achieving superior performance in predicting RNA-protein interactions. For peptide-binding residues, several approaches have been formulated. Taherzadeh et al. developed SPRINT-Seq ^21^, a Support Vector Machine-based method that uses sequence-based features to predict peptide-binding residues. Zhao et al. (2018) constructed a consensus-driven method named PepBind ^22^, pioneering the inclusion of intrinsic disorder into feature design, as research indicates a strong correlation between protein-peptide binding and intrinsic disorder. Moreover, Wardah et al. presented Visual ^23^, a CNN-based technique that transcodes protein sequences into image-like forms for predicting peptide-binding residues in proteins. In the realm of multi-type prediction methods, which are capable of predicting the interacting residues between proteins and different molecules simultaneously, several methods have been proposed. Yan et al. presented a novel sequence-based method, DRNApred ^24^, developed to accurately predict and discriminate between DNA– and RNA-binding residues by leveraging a unique dataset with both DNA– and RNA-binding proteins. Furthermore, Wang et al. proposed iDRNA-ITF ^25^, a sequence-based method integrating functional properties of residues through an induction and transfer framework to enhance the identification of DNA– and RNA-binding residues in protein sequences. Additionally, Zhang et al. developed an innovative deep multi-task architecture, DeepDISOBind ^26^, designed to accurately predict DNA-, RNA– and protein-binding intrinsically disordered regions (IDRs) from protein sequences by employing an information-rich sequence profile.

Despite the development and achievements of computational methods in recent years, they encounter certain issues that constrain their applicability for large-scale, high-throughput predictions. First, many of the existing methods make predictions based on structural information. Due to the lack of knowledge about most protein-ligand complex structures, current structure-dependent prediction methods are significantly reliant on structural data derived from external computational tools. This dependence can readily cause a decrease in prediction effectiveness and introduce extraneous noise, thereby negatively influencing the performance of the prediction. Second, many sequence-based methods rely on biological feature engineering, which poses similar challenges for them. For instance, the evolutionary information frequently utilized by these methods is derived from position-specific scoring matrices (PSSM). These matrices are generated through sequence alignment between the query proteins and extensive protein databases. The process of extracting and integrating various biological features involves substantial manual curation and domain expertise, resulting in high costs and time investments. Third, single-type prediction methods, can only predict interacting residues between proteins and a specific molecule, which limits their practical applicability. While multi-type prediction methods aim to overcome this limitation by predicting interactions with multiple types of molecules simultaneously, they still have their own set of constraints. One notable limitation of multi-type prediction methods is that they inherently rely on different neural network architectures for each type of interaction. This means that for predicting interactions with DNA, RNA, peptides, and other molecules, separate models or network structures need to be developed, trained, and maintained. This increases the complexity and computational resources required for training and deploying these models. Another limitation of multi-type prediction methods is the extensive use of biological feature engineering, leading to considerable expenses and time commitments.

To address these issues, we have established a new ligand-type-aware contrastive learning framework, referred to as MucLiPred, for the prediction of multiple protein-ligand binding residues. The innovative aspects of our method can be succinctly encapsulated as follows:

1) Setting it apart from conventional methods that rely on other information predicted by third-party tools, MucLiPred is a purely sequence-based prediction method that utilizes only protein sequences for model training and prediction, thereby enhancing the prediction speed and computational efficiency.
2) We introduce a new module called residue-level contrastive learning to tackle the issue of data imbalance. It is capable of adaptively learning more discriminative and high-quality representations of the binding residues, fully utilizing the majority of samples in the imbalanced dataset.
3) We proposed a type-level contrastive learning module to focus on the ligand type information of the input sequences, which allows the model to pay attention to the category information of the sequences, forming different feature representations for different categories, thereby enabling the model to predict the binding residues of different types of molecules simultaneously.
4) We utilized type-level contrastive learning, enabling the model to predict the binding residues of different types of molecules while using the same set of parameters. This overcomes the high time and computational resource expenses associated with multi-type prediction methods. Moreover, during the process of predicting different types, the model can obtain some auxiliary information through shared layers, thereby improving the model’s generalization ability.

For instance, if the model learned the binding residues information of a certain protein sequence and DNA during the training process, it could utilize the previously learned information related to that sequence to better predict the binding residues with RNA during the prediction process. For two different protein sequences, they may also have similar sequence fragments, thereby enhancing each other’s prediction effectiveness.

## 2 Materials and methods

### 2.1 Datasets

For academic research, we have opted for two frequently employed reference datasets to impartially assess and contrast our proposed approach against established methods. For the sake of convenience in our discourse, we have designated the aforementioned datasets as **Dataset 1** and **Dataset 2**, correspondingly.

**Description of Dataset 1**. The dataset was proposed by the study of a deep learning-based method called DBPred ^17^ for the prediction of DNA interacting residues in a protein. It contains 692 protein-DNA complexes with a total of 16601 binding residues (positive) and 308414 non-binding residues (negative). The training set of **Dataset 1** contains 646 proteins with 15636 binding residues and 298503 non-binding residues, while the testing set of **Dataset 1** comprises 46 proteins with 965 binding residues and 9911 non-binding residues.

**Description of Dataset 2**. The dataset is sourced from the research conducted on Pprint2 ^27^, a convolutional neural network-based method for the purpose of predicting protein-RNA interacting residues. It comprises a collection of 706 protein-RNA complexes, encompassing a total of 25525 binding residues (positive) and 216228 non-binding residues (negative). The training set of **Dataset 2** consists of 545 proteins, encompassing 18559 binding residues and 171879 non-binding residues. Furthermore, the testing set of **Dataset 2** comprises 161 proteins, containing 6966 binding residues and 44349 non-binding residues.

Figure 1A illustrates the pre-processing steps we conducted on the datasets. It is important to highlight that both datasets underwent a similar pre-processing procedure in order to facilitate the training of reliable predictive models. We performed truncation on the sequence data in the datasets, restricting their lengths to fall between 200 and 500. By applying truncation to the datasets, it becomes possible to confine the lengths of sequences to a reasonable range. This, in turn, alleviates the computational burden and memory limitations, leading to improved training efficiency of the model. Moreover, longer sequences often contain redundant information or noise that can impede the model’s training and generalization capabilities. Through truncation, selected segments of the excessively long sequences are eliminated, enabling the model to concentrate on crucial information, effectively controlling model complexity, and enhancing overall performance. After performing truncation operations on Dataset 1 and Dataset 2 separately, we introduced ligand type information into the datasets. For the dataset used for predicting protein-DNA binding (Dataset 1), we included the conditional information “DNA binding” for each data entry. Similarly, for the dataset used for predicting protein-RNA binding (Dataset 2), we also included the conditional information “RNA binding” for each data entry. Following the introduction of ligand type information, we merged these two datasets to obtain the final dataset named **Merged Dataset** used for training. It is worth noting that, for improved training performance, we ensured consistency in the number of sequences included in Dataset 1 and Dataset 2. In essence, the balanced treatment of different ligand types in our preprocessing steps ensures a robust and unbiased learning environment for our model. This paves the way for effective contrastive learning, where the model benefits from the nuanced understanding of distinct ligand-specific protein binding characteristics, thereby enabling accurate and diverse protein-ligand binding predictions.

**Figure. 1.**
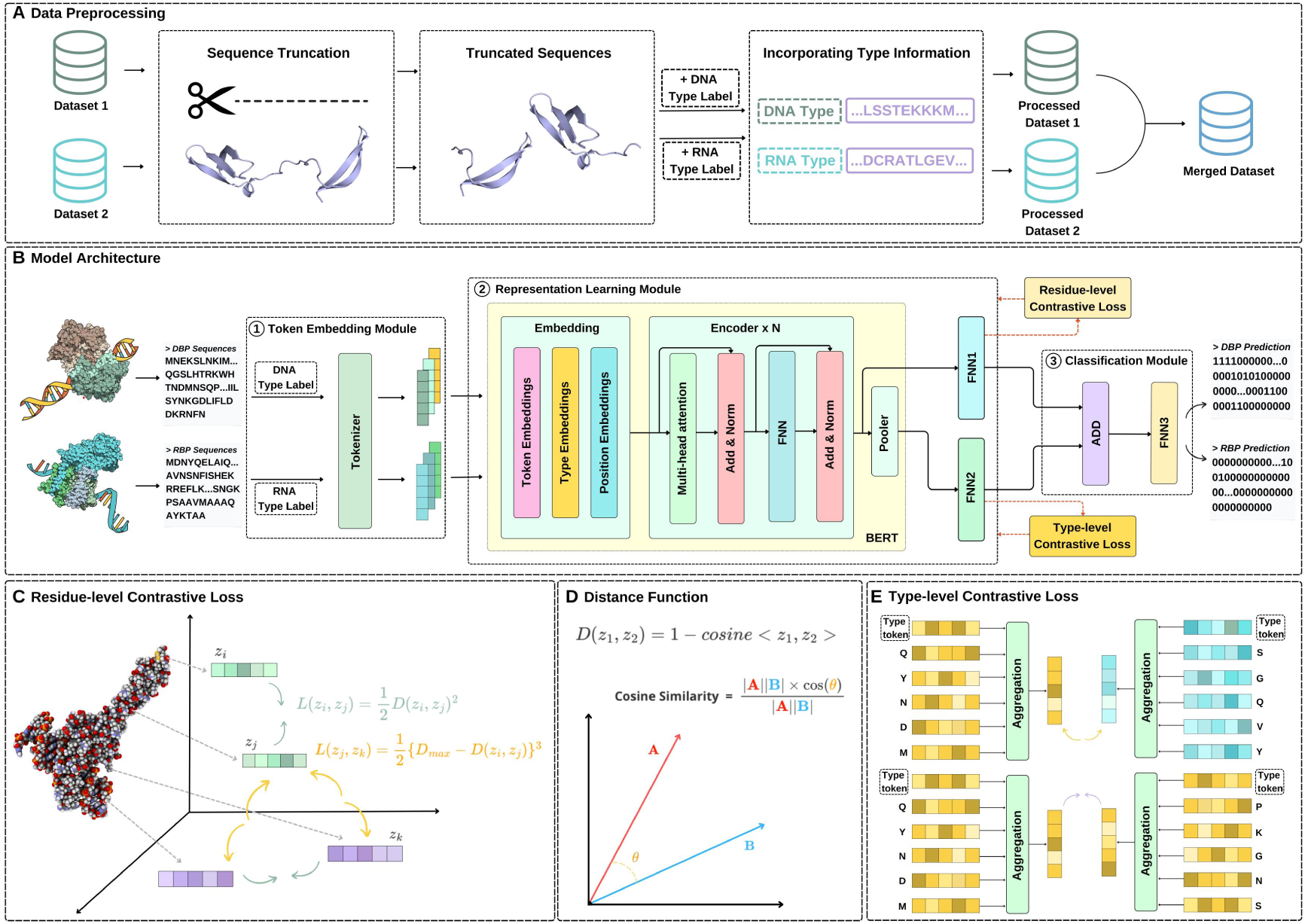
The workflow and framework of the proposed MucLiPred. (**A**) Dataset processing. Sequences from Dataset 1 and 2 were truncated to lengths between 200 and 500, and annotated with ligand type information. The processed datasets were then merged to form a comprehensive training dataset called Merged Dataset. **(B)** Architecture of deep learning network in MucLiPred. The architecture consists of three modules: (i) Token embedding module: This module transforms raw protein sequences and ligand type information into pre-trained embedding vectors, (ii) Representation learning module: This module leverages a pre-trained BERT model to encode the embedding matrix into high-dimensional representation vectors and optimizes the contrastive loss at both residue and type levels, and (iii) Classification module: This module merges dimensionality-reduced residue and type-level features, transforming them into final binding residue predictions through a Feed-Forward Neural Network. **(C)** The process of calculating residue-level contrastive loss. The model calculates and optimizes the residue-level contrastive loss to accentuate the distinctions between different classes of residues, essentially those that are binding and those that are not. **(D)** The definition of distance function. The distance function quantifies the distance between the representations of a pair of residues or ligand types, which is used in the calculation of contrastive loss. **(E)** The process of calculating type-level contrastive loss. The model calculates and optimizes the type-level contrastive loss to further differentiate the ligand-binding characteristics of various ligand types. This process ensures that samples from the same ligand type are closer in the representation space, while those from different ligand types are farther apart.

### 2.2 Architecture of the proposed model

Figure 1B displays a schematic representation of the model architecture utilized in this research, comprising a few modules: (1) Token Embedding Module, (2) Representation Learning Module, and (3) Classification Module.

Within the token embedding module, our model takes the query protein sequence with ligand type information as input. In this module, each amino acid present in the sequence is transformed into a pre-trained embedding vector, effectively encoding the sequence. Consequently, the sequence is converted into an embedding matrix.

In the Representation learning module, the obtained embedding matrix undergoes additional encoding using a pre-trained BERT model, resulting in the generation of a representation vector with high dimensionality. This approach effectively mitigates the challenge of long-distance dependency in protein sequences when identifying ligand-binding residues, thanks to the utilization of multi-head attention mechanism. We calculate and optimize the residue-level contrastive loss between any two training samples in the training set, and thus obtain more discriminative representations of the binding (non binding) residues. Additionally, we apply the type-level contrastive loss to further differentiate the ligand-binding characteristics of various ligand types. Leveraging the power of contrastive learning at the type level, our model learns to distinguish between distinct classes of ligands, thereby refining the representation of different ligand-binding behaviors.

In the Classification Module, we merge the output from the BERT Encoder and the output from the Pooler layer that contains ligand type information using the addition operation.

Subsequently, we pass this combined output through several layers of a fully connected network to obtain the final prediction result.

### 2.3 Token embedding module

The initial step involves translating the raw protein sequence into a digital sequence, employing a predefined vocabulary dictionary. In this dictionary, each amino acid in the sequence corresponds to a word in a sentence and is mapped to a specific numerical value. Consequently, the raw protein sequence is encoded as a vector of numerical values. The BERT model employs three distinct types of embeddings to capture various nuances in the input data: Token Embeddings, Segment Embeddings, and Position Embeddings ^28^.

Token Embeddings are utilized to transform each token (amino acid in this context) into a representative vector carrying its contextual semantics. This process is similar to mapping each word in a sentence to a multi-dimensional space, where semantically similar tokens are closer to each other. Position Embeddings play a pivotal role in preserving the order within the sequence. In the context of natural language, the arrangement of words significantly impacts the overall meaning of a sentence. Similarly, the order of amino acids within a protein sequence is crucial in determining its function. Hence, Position Embeddings assign a unique vector to each position in the sequence, enabling the model to learn and maintain the inherent sequential information. Segment Embeddings, in the original BERT model, are designed to differentiate and identify two different sentences or segments. They are typically employed for sentence pair classification tasks where the input consists of two sentences, as in question-answering systems and natural language inference tasks. The introduction of Segment Embeddings allows the model to distinguish and understand the different contexts of these two sequences, assigning a unique embedding for each.

Subsequently, we utilized the Segment Embeddings of the BERT model to incorporate ligand type information into our input, distinguishing between different ligand types in protein sequences ^29^. This allowed our model to learn the relationships among sequences of different types. For clarity, we renamed Segment Embedding as Type Embedding, reflecting its role in encoding and distinguishing protein sequence types more appropriately.

### 2.4 Representation learning module

#### 2.4.1 Encoding process of the module

BERT, a bidirectional language representation model based on the transformer architecture ^30^, has become a staple in various NLP tasks due to its powerful language understanding capabilities. This success can be attributed to the extensive pre-training that BERT undergoes, often on a substantial corpus related to the target domain.

In our research, we leveraged the pre-trained ProtBert-Big Fantastic Database (BFD) ^31^, a BERT variant that has been specifically adapted for protein sequences. This model was pre-trained using a large corpus of unlabeled protein data from the BFD dataset ^32^, a compendium of 2.1 billion protein sequences. Unlike the typical BERT pre-training process, this model was trained exclusively using the masked language model technique, given the absence of semantic logic among protein sequences. The fundamental building block of the BERT model is the Encoder block, which comprises multiple components including a multi-head attention mechanism, a feed-forward neural network (FNN), and the residual connection technique. The multi-head attention mechanism consists of several individual self-attention modules, enabling the model to learn diverse contextual representations of protein sequences from multiple perspectives.

The multi-head attention mechanism of the BERT model allows the model to focus on different parts of the input sequence for each attention head, effectively capturing various aspects of the sequence. The multi-head attention is based on the self-attention module and the self-attention mechanism is described as follows:

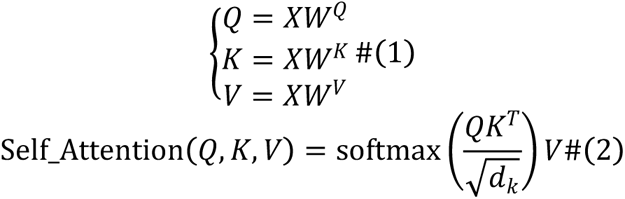

where Q, K, and V denote the query, key, and value vectors and are formulated as Q = XW^Q, K = XW^K, and V = XW^V. The matrices W^Q, W^K, and W^V serve as the respective weight matrices for Q, K, and V.

The multi-head attention mechanism essentially employs several parallel self-attention processes, known as “heads”. Each head computes its own attention scores using the query (Q), key (K), and value (V) matrices derived from the input X. This setup allows different heads to capture various types of sequence relationships. Post computation, the outputs from all heads are concatenated and linearly transformed to produce the final representation. This amalgamation enables the model to gain a richer and more comprehensive understanding of the sequence by leveraging insights from multiple attention perspectives.

After the encoding process of the BERT model, we obtain y_r, which is the output of the last Encoder block. Subsequently, y_r serves as the input to the Pooler block of the BERT model, generating the output y_t. The Pooler block is a dense layer that takes the first token’s output (the “[CLS]” token) from the final Encoder block as input. It plays a crucial role in sentence-level classification tasks, providing an aggregated representation of the entire input sequence.

To manage the high dimensionality of y_r and y_t, we employed two Feed-Forward Neural Network (FNN) blocks for dimensionality reduction. FNN1 targets y_r, and FNN2 is dedicated to y_t. By incorporating this approach, we enable the model to extract a more concise representation of the amino acids in the input sequence while maintaining its unique features. Consequently, this significantly enhances the contrastive learning efficiency and overall performance of our model.

#### 2.4.2 Residue-level contrastive learning

The surge in available augmented data, bolstered by various text data augmentation techniques, has fostered the widespread adoption of contrastive learning in natural language processing (NLP). This learning method, which thrives on distinguishing between positive and negative samples, leverages the abundant augmented data to improve the learning environment ^33^. This growth in usage has led to the emergence of innovative contrastive learning frameworks in NLP, enriching the field with diverse methodologies and perspectives ^34^. Due to the task characteristics, most existing contrastive learning frameworks primarily rely on unsupervised learning and lack explicit constraints on negative samples ^35^. In this work, we introduce a novel residue-level contrastive learning module that operates within the framework of supervised learning.

Our module aims to ensure that representations of inputs from the same class are mapped to neighboring points in the representation space, while inputs from different classes are mapped farther apart. Specifically, considering the variation in protein sequence lengths and to maintain consistency, we collect a batch-size of representation matrices from the encoder module without padding sequences to the same length. This approach allows us to gather an adequate amount of residue-level data for an effective contrastive learning ^36^. As a subsequent step, we aim to ensure that samples from the same class possess similar representations, while samples from different classes have dissimilar representations.

To achieve this, we introduce a residue-level contrastive loss function, denoted as *L_rc_*, within our framework. This loss function is designed to address the challenge posed by imbalanced datasets, promoting the learning of discriminative representations. For a pair of residue representations z_1, z_2 in a batch, the loss is defined as follows:

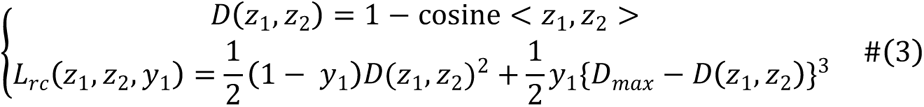

The distance between the representations of a pair of residues, denoted as z_1 and z_2, can be quantified using the D(z1,z2). The maximum value of D(z1, z2), denoted as D_max, is set to two in this case. If a pair of residues belong to different classes, indicating that one residue is binding while the other is not, the corresponding label y_1 is assigned a value of 1. Conversely, if the pair of residues belong to the same class, the label y _1 is set to 0. Importantly, we enhance the power of the different-class pair to three in our model, indirectly prompting the system to dedicate increased attention to the minority class, following the approach taken in PepBCL ^36^.

#### 2.4.3 Type-level contrastive learning

While residue-level contrastive learning focuses on creating discriminative representations at the micro-level (residue), it is equally important to address this task at a macro-level (ligand type or protein). This has led to the development of our type-level contrastive learning module, which operates in the broader context of protein types.

Just as the residue-level module aims to differentiate between positive and negative samples at the residue level, the type-level module aims to ensure that samples from the same ligand type are closer in the representation space, while those from different ligand types are further apart. The core of this approach is the collection of a batch size of representation vectors from the FNN2 module, which reduces the dimensionality of the pooled sequence representation, ***y_t***. This procedure allows us to gather sufficient data at the protein sequence level for effective contrastive learning.

To augment the learning process, we incorporate an additional contrastive loss function into our framework, designated as *L_tc_*. Analogous to its residue-level counterpart, this loss function aims to bolster the differentiation and learning of unique ligand type representations, concurrently assimilating distinct ligand binding residue patterns.

In our model, for a pair of ligand type representations (i.e. the output ***y_t*** from the Pooler block) ***t_1***, ***t_2*** in a batch, the loss is defined as follows:

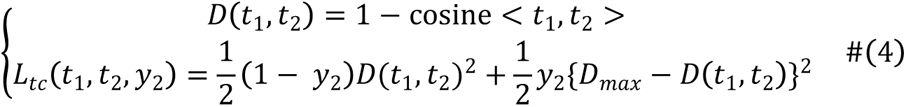

If a pair of representations belongs to the same ligand type, the corresponding label ***y_2*** is assigned a value of 0. On the contrary, if the representations pertain to different ligand types, the label ***y_2*** is set to 1.

Through this approach, our model can effectively learn discriminative representations both at the residue level and the ligand type level, thus providing a more comprehensive understanding of protein structures. This dual-contrastive learning mechanism is key to our model’s robustness and adaptability.

### 2.5 Classification module

Initially, our model generates a residue-level representation vector, ***y_r***, by conducting a feature vector transformation on the raw protein sequence, ***x***. This transformation process is facilitated by the Encoder. Simultaneously, a type-level representation vector, ***y_t***, is produced by transforming the ligand type label information through the Pooler block. Subsequently, we employ FNN1 and FNN2 to perform dimensionality reduction on ***y_r*** and ***y_t*** respectively. Following this, an addition operation is performed on the dimensionality-reduced residue-level representation vector ***FNN1(y_r)*** and the type-level representation vector ***FNN2(y_t)***. This operation effectively fuses and enhances their respective feature information, yielding an integrated representation vector, ***z***. Finally, a Feed-Forward Neural Network called FNN3 processes the integrated representation vector ***z***, transforming it into a residue-level category output, ***y_p***. This step effectively translates the rich information encapsulated within ***z*** into a format that the model can utilize for residue-level classification.

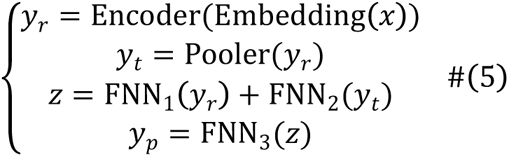

In order to enhance the prediction performance, the output module is trained using the cross-entropy loss function ***L_ce***.

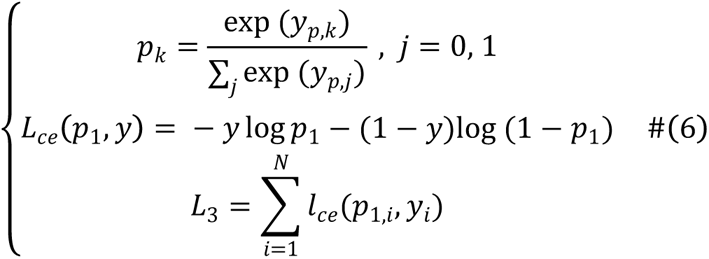

In this context, ***k*** equals either 0 or 1, each corresponding to the class of non-binding residues or binding residues respectively, and ***p_k*** represents the likelihood of a residue being classified as category ***k***. N is the total count of residues in a batch, ***y_i*** stands for the label associated with the residue ***i***, and ***L_3*** signifies the cross-entropy loss incurred for a batch.

In order to prevent the backpropagation of loss ***L_3*** from interfering with the representation learning module and to circumvent the gradient diminishing issues, we opt to isolate the optimization processes of the representation learning module and the classification module. Specifically, during the training phase of the classification module, we keep the parameters within the representation learning module constant. Therefore, the loss function of our model can be formulated as follows:

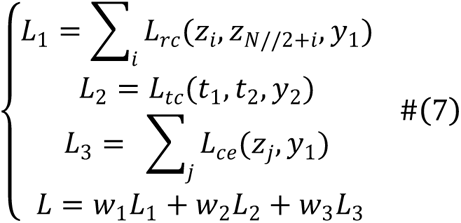

where ***w_1***, ***w_2,*** and ***w_3*** serve as weighting coefficients to modulate the relative influence of each constituent loss function. These weights facilitate the balanced integration of the distinct loss components, ensuring that no single loss dominates the learning process and enabling fine-tuned control over the model’s learning dynamics.

### 2.6 Evaluation metrics

In this research, the independent datasets exhibit a significant imbalance between positive and negative samples. To comprehensively assess the performance of our proposed method, we employ five widely-used metrics: ACC, Precision, Matthews correlation coefficient (MCC), F1 Score, and Area Under the ROC Curve(AUC). The calculation formulas are as follows:

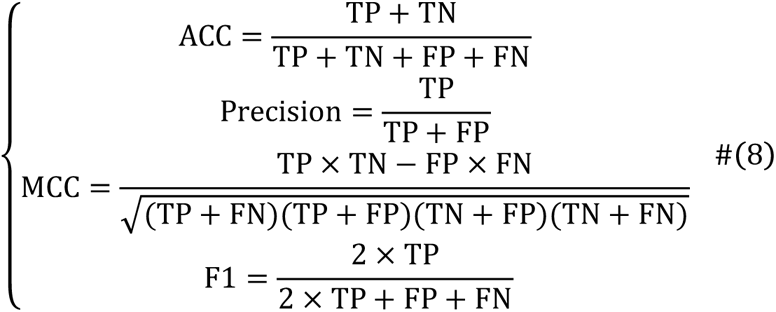

In the formulas provided above, TP (true positive) and TN (true negative) represent the number of correctly predicted binding residues and non-binding residues, respectively. Conversely, FP (false positive) and FN (false negative) denote the number of binding residues and non-binding residues that were incorrectly predicted. ACC (Accuracy) measures the overall correctness of the model’s predictions, taking into account both correct and incorrect predictions. MCC (Matthews Correlation Coefficient) is a balanced measure that considers both the prediction ability for binding residues and non-binding residues, making it a highly suitable metric for imbalanced datasets. Precision assesses the accuracy of residues identified as binding in predictions. The F1 score is the harmonic mean of Precision and Recall, providing a balanced metric that evaluates the model’s performance in terms of both false positives and false negatives. AUC (Area Under the Receiver Operating Characteristic Curve) measures the performance of the binary classification model at all classification thresholds. This metric provides a comprehensive view of the model’s performance, as it is not tied to a specific threshold. These measures collectively offer a comprehensive assessment of our model’s performance, taking into account various aspects of binary classification tasks and catering to the nature of imbalanced datasets.

## 3 Results

### 3.1 Comparison with existing methods

To validate the effectiveness of our newly developed method MucLiPred, it is crucial to conduct a performance comparison with existing methods. For the prediction of protein-DNA binding residues, we conducted a comparative analysis with five existing methods: DRNAPred, DNAPred, ProNA2020, NCBRPred, and DBPred. It is worth mentioning that among the five methods, DRNAPred, ProNA2020, and NCBRPred are multi-type predictors, which means they are capable of predicting binding residues for both DNA and RNA. On the other hand, the remaining methods are single-type predictors, focusing solely on DNA binding residues. We conducted comparative experiments on the testing sets of Dataset 1. The results of the comparisons are presented in **Table 2**. It should be noted that the source codes for some of the compared methods are not available. As a result, the results for these methods on the two datasets are taken directly from their studies. As indicated in Table 2, our proposed MucLiPred exhibits notably superior performance compared to the single-type prediction methods (i.e. DNAPred and DBPred) on the testing sets of Dataset 1. More specifically, MucLiPred demonstrates an ACC of 88.36%, an AUC of 0.86, and an MCC of 0.44, representing a relative improvement of 10.64%, 7.00%, and 12.00% over DBPred, respectively. Simultaneously, our proposed MucLiPred demonstrates significantly enhanced performance when compared to the multi-type prediction methods (i.e. DRNAPred, NCBRPred, and DBPred) on the testing sets of Dataset 1. To provide more specific details, our proposed method MucLiPred demonstrates a significant improvement over NCBRPred, with relative improvements of 20.90%, 15.00%, and 23.00%, respectively. Furthermore, the bar chart presented in Figure 2A highlights the performance of different methods, including MucLiPred, in terms of ACC, AUC, and MCC on the testing set of Dataset1. The results demonstrate that MucLiPred consistently outperforms other existing methods across all three metrics, achieving the highest AUC, AUC, and MCC.

**Table 1.** Summary of datasets.

| Dataset | Dataset 1 |  | Dataset 2 |  |
| --- | --- | --- | --- | --- |
|  | training | testing | training | testing |
| Number of proteins | 646 | 46 | 545 | 161 |
| Number of residues | 314139 | 10876 | 190438 | 51315 |
| Number of binding residues | 15636 | 965 | 18559 | 6966 |
| Number of non-binding residues | 298503 | 9911 | 171879 | 44349 |

**Figure. 2.**
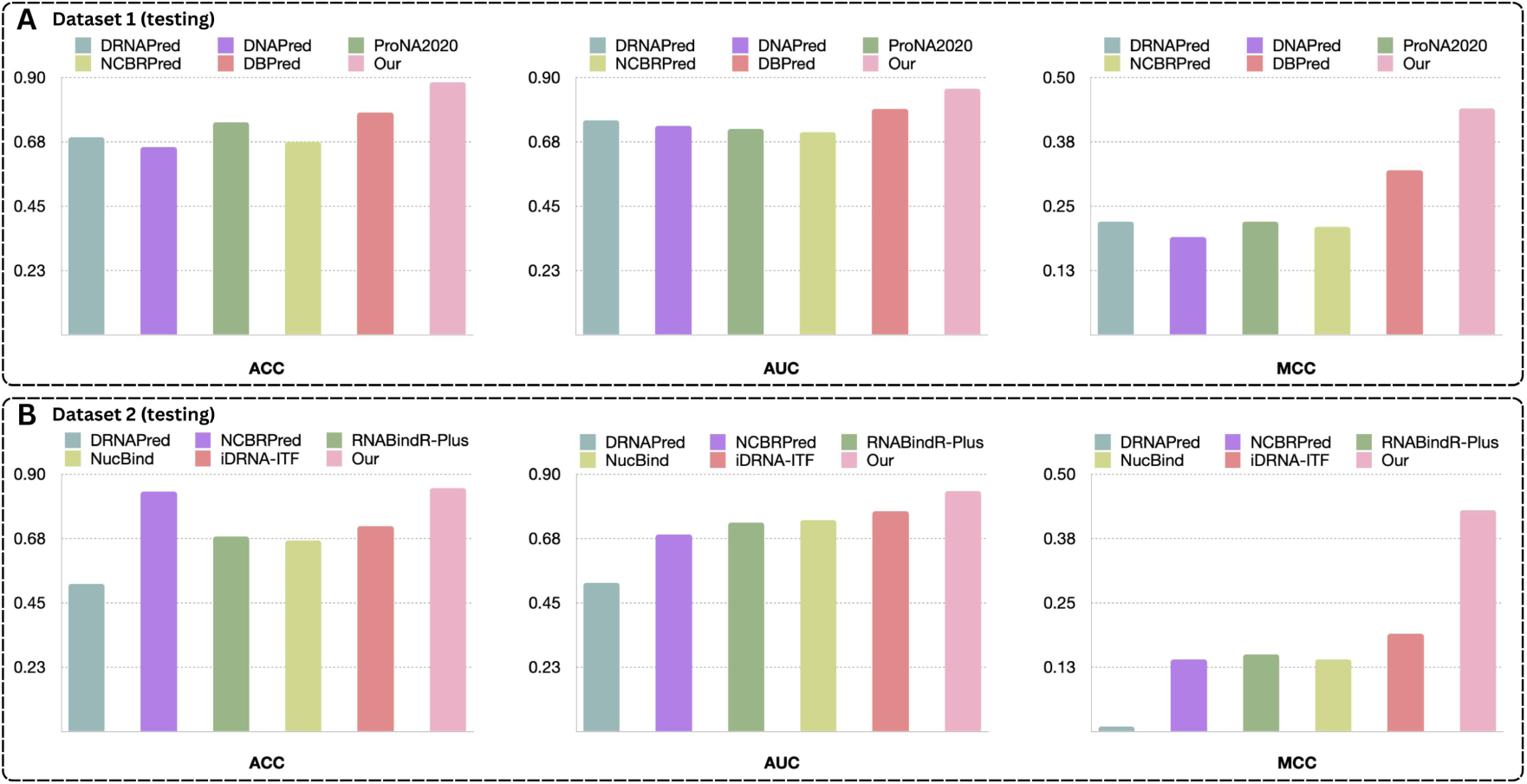
Comparative performance of our proposed MucLiPred method versus other established methods on benchmark datasets. (**A**) shows performance comparison on Dataset 1. **(B)** shows performance comparison on Dataset 2.

**Table 2.** Comparison of the proposed MucLiPred and other methods on the testing sets of Dataset 1.

| Model | ACC (%) | AUC | MCC |
| --- | --- | --- | --- |
| DRNAPred* | 69.06 | 0.75 | 0.22 |
| DNAPred | 65.64 | 0.73 | 0.19 |
| ProNA2020* | 74.31 | 0.72 | 0.22 |
| NCBRPred* | 67.46 | 0.71 | 0.21 |
| DBPred | 77.72 | 0.79 | 0.32 |
| Our | **88.36** | **0.86** | **0.44** |

*Note:* Metrics for other methodologies were sourced from their respective publications. Methods marked with an asterisk (\*) are multi-type prediction methods. The bold typeface signifies the top performance within each metric.

In the assessment of protein-RNA binding residues prediction, we conducted a comparative analysis with five existing methods, namely DRNAPred, NCBRPred, RNABindR-Plus, NucBind, and iDRNA-ITF. It should be noted that among the five methods examined, DRNAPred, NCBRPred, and iDRNA-ITF are considered multi-type predictors, capable of predicting binding residues for both DNA and RNA. Conversely, the remaining methods focus exclusively on RNA binding residues, making them single-type predictors. We performed comparative experiments on the testing sets of Dataset 2 and presented the corresponding results in **Table 3**. As observed in **Table 3**, our proposed MucLiPred shows significantly superior performance compared to the single-type prediction methods (i.e. RNABindR-Plus and NucBind) on the testing sets of Dataset 2. Specifically, MucLiPred achieves an accuracy of 85.16%, an AUC of 0.84, and an MCC of 0.43, resulting in a relative improvement of 18.29%, 10.00%, and 29.00% over NucBind, respectively. Furthermore, our proposed MucLiPred demonstrates remarkable performance improvements when compared to the multi-type prediction methods (i.e. DRNAPred, NCBRPred, and iDRNA-ITF) on the testing sets of Dataset 2. To provide more specific details, MucLiPred outperforms iDRNA-ITF with substantial relative improvements of 13.33%, 7.00%, and 24.00%, respectively. The bar chart presented in Figure 2B illustrates the performance of different methods in terms of ACC, AUC, and MCC on the testing set of Dataset 2. The results consistently indicate that MucLiPred outperforms other existing methods across all three metrics, achieving the highest ACC, AUC, and MCC.

**Table 3.** Comparison of the proposed MucLiPred and other methods on the testing sets of Dataset 2.

| Model | ACC(%) | AUC | MCC |
| --- | --- | --- | --- |
| DRNAPred* | 51.59 | 0.52 | 0.01 |
| NCBRPred* | 83.95 | 0.69 | 0.14 |
| RNABindR-Plus | 68.31 | 0.73 | 0.15 |
| NucBind* | 66.87 | 0.74 | 0.14 |
| iDRNA-ITF* | 71.83 | 0.77 | 0.19 |
| Our | **85.16** | **0.84** | **0.43** |

*Note:* Metrics for other methodologies were sourced from their respective publications. Methods marked with an asterisk (\*) are multi-type prediction methods. The bold typeface signifies the top performance within each metric.

The multi-type prediction methods DRNAPred and NCBRPred, listed in **Tables 1** and **Table 2**, exhibit better performance in predicting DNA binding residues than RNA ones, as evidenced by superior ACC, AUC, and MCC scores on Dataset 1(Testing) compared to Dataset 2(Testing). In other words, for these two multi-type prediction methods, their performance in predicting DNA binding residues is superior to predicting RNA binding residues. One possible explanation for this phenomenon is that RNA binding residues are generally less abundant compared to DNA binding residues. Even when using comparable size datasets for training the model, it becomes more challenging for the model to identify the binding patterns and distribution characteristics of RNA binding residues. As a result, the predictive performance of the model for RNA binding residues is inferior to that for DNA binding residues. However, our model MucLiPred avoids the issue of inconsistent predictive performance for DNA and RNA binding residues. When evaluated on Dataset 1(Testing), our model achieved AUC and MCC scores of 0.86 and 0.44, respectively. Similarly, when assessed on Dataset 2(Testing), our model attained AUC and MCC scores of 0.84 and 0.43, respectively. Crucially, our model utilized a merged dataset during the training process, allowing it to simultaneously learn the common characteristics and distinctions in the distribution of DNA and RNA binding residues. This Merged dataset, to some extent, can be considered as an augmentation of the learning samples for RNA binding residues. As a result, the model benefits not only from the learned distribution information of RNA binding residues but also from the distribution patterns of DNA binding residues as auxiliary information during the prediction of RNA binding residues. This approach enhances the model’s predictive performance and mitigates the issue of performance inconsistencies.

### 3.2 Exploration of the optimal model architecture

In order to achieve optimal performance for our model, we continuously adjusted and explored various components of the model. The specific adjustments included:

1) Selection of feature vectors used in the type-level contrastive learning process.
2) The approach for incorporating type information in the classification module.
3) Determination of the location for dimensionality reduction of feature representations.

#### 3.2.1 Selection of feature vectors

In the process of contrastive learning, the selection of appropriate feature vectors is of paramount importance. When properly chosen, feature vectors facilitate a more efficient measure of similarity and dissimilarity among data points, thereby fostering the learning of distinct data representations. Furthermore, the appropriateness of the feature vectors can greatly influence the efficiency of the learning algorithm. Lastly, a good choice of feature vectors not only delivers excellent performance on training data but also generalizes well to unseen. In the process of type-level contrastive learning, we opted for two distinct approaches to select feature vectors. Firstly, we utilized the mean pooling operation on the output of the Encoder to obtain a vector representation. Secondly, we incorporated the output from the Pooler layer as the feature vector for conducting contrastive learning. Additionally, we also explored a configuration in our model that excluded type-level contrastive learning as a point of reference.

We presented the performance of the model under different approaches in Figure 3 using bar charts. It is evident that the group utilizing the output from the Pooler layer as feature vectors achieved the best results across AUC, F1, and MCC metrics. The group utilizing mean pooling, on the other hand, exhibited the poorest performance across all three metrics. The discrepancy in performance between the two groups can be attributed to the way they handle the sequence information. When applying mean pooling, the model takes an average of all vectors in the sequence, yielding a single vector representation. While this method is computationally simple and efficient, it has several significant drawbacks. Firstly, mean pooling disregards the ordering of elements in the sequence, failing to capture the order information, which in turn can severely affect the learning outcome. Secondly, mean pooling assigns equal weights to all elements, thereby neglecting the varying contribution of different elements to the overall sequence meaning. Lastly, mean pooling could decrease the distinctiveness between different sequences, as it may map diverse sequences to similar vectors. Conversely, the group utilizing the output from the Pooler layer exhibited superior performance. The Pooler layer, positioned following the last Transformer encoder, processes the final hidden state of the initial sequence token to create a summary representation that captures vital global semantic information. This is especially important for contrastive learning. Additionally, the [CLS] token in BERT, which provides a weighted contextual summary of the entire sequence, captures rich linguistic nuances, thereby offering a more nuanced sequence representation than simpler pooling methods and enhancing contrastive learning. As a result, utilizing Pooler output proves to be an efficient strategy for high-quality sequence representations in contrastive learning.

**Figure. 3.**
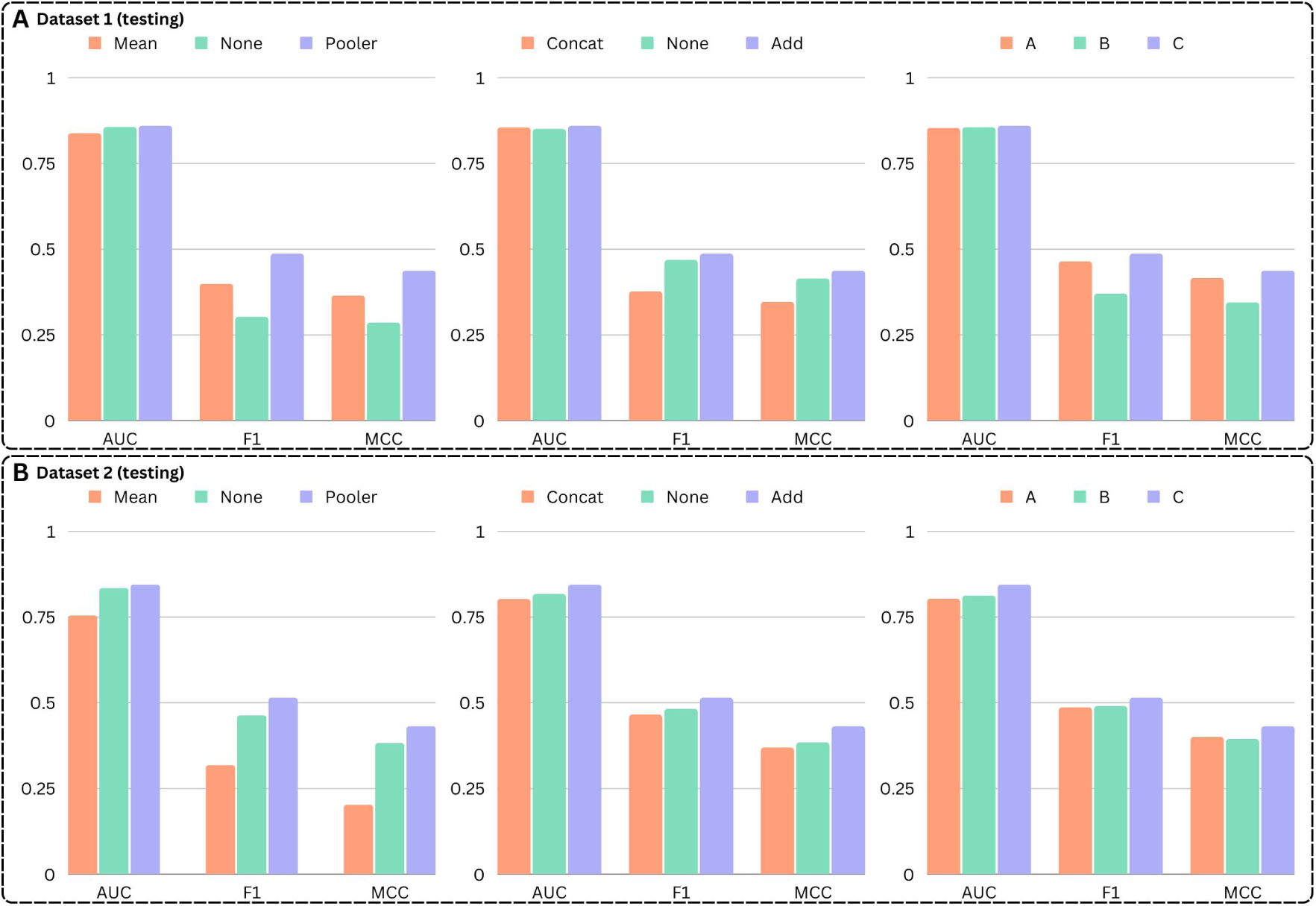
Comparison of model performance under different architectures. It showcases the impact of different approaches for selecting feature vectors, incorporating type information, and applying dimensionality reduction on the model’s performance. **(A-B)** illustrate the performance of various model configurations using bar charts across AUC, F1, and MCC metrics on Dataset 1 and 2, respectively. The detailed raw data corresponding to these figures can be found in **Supplementary Tables 1** and **2**.

#### 3.2.2 The approach for incorporating type information

In deep learning, information fusion is to improve the performance and representation capabilities of the model by combining the contributions of multiple information sources. In the classification module, we incorporate the ligand type information of the sequence into the output of the BERT encoder. This allows the classification layer to pay more attention to the category information of the sequence, facilitating better prediction and classification.

We selected both addition (add) and concatenation (concat) as the methods for information fusion, and we included a group that did not perform any information fusion as a control. In Figure 3, it can be observed that the group utilizing the addition method to incorporate ligand type information achieved the best results across all metrics. However, the group employing the concatenation method obtained the poorest performance indicators. The differences in performance can be attributed to the intrinsic properties of the addition and concatenation methods for information fusion.

The addition is an element-wise operation, and it combines the features by superimposing the individual values from both feature vectors. This implies that the features from both the BERT encoder and the type information are blended into the same space, preserving the original feature dimensionality. Through element-wise addition, critical insights from both inputs are retained and merged, enhancing the prominence of influential features within the inputs. On the other hand, concatenation preserves all original features as it directly combines the feature vectors end-to-end. The increase in feature dimensionality can lead to higher computational requirements and could potentially increase the risk of overfitting, as the model has a larger feature space to fit the data. Additionally, concatenation maintains the features in separate spaces, which could make it harder for the model to establish relationships between the features from the Encoder and the type information. This separation can result in suboptimal performance when the interplay between features is critical for the prediction task. In summary, the superior performance of the addition-based fusion, in this case, could be attributed to its capability to blend the feature spaces and establish direct interactions between features, thereby effectively leveraging the combined information for prediction. On the contrary, the increased feature dimensionality and potential hindrance to feature interactions might have led to the poor performance of concatenation-based fusion.

#### 3.2.3 Determination of the location for dimensionality reduction

Dimensionality selection is a critical aspect to consider in contrastive learning. It directly impacts the performance and representation capability of the model. Based on whether dimension reduction was applied and where it was applied, we categorized the model structures into three types: A) No dimension reduction applied to the feature vectors used in either residue-level or type-level contrastive learning; B) Dimension reduction applied to the feature vectors used in residue-level contrastive learning but not in type-level contrastive learning; C) Dimension reduction applied to the feature vectors used in both residue-level and type-level contrastive learning. Their respective structures are illustrated in Figure 4.

**Figure. 4.**
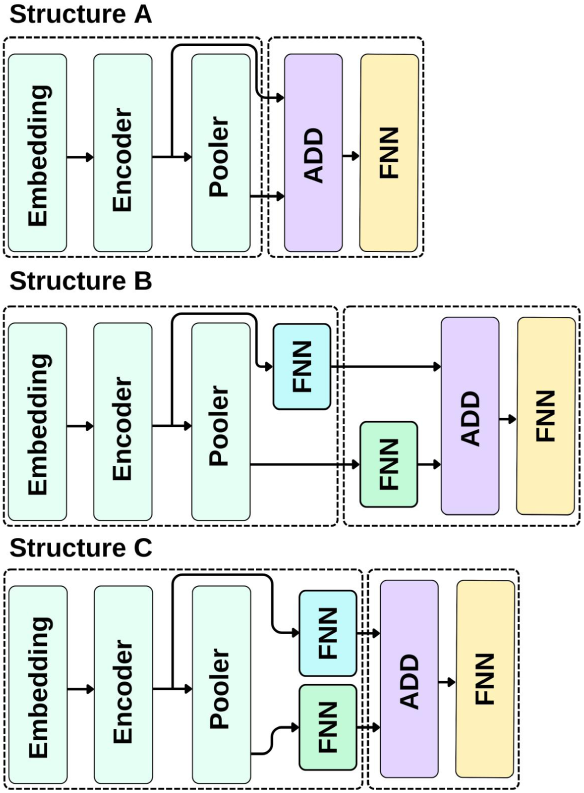
Three variations of our model structures. (**A**) Structure A: No dimension reduction. **(B)** Structure B: Dimension reduction in residue-level only. **(C)** Structure C: Dimension reduction in both residue-level and type-level.

Comparing structure A with structure C, as depicted in Figure 3, the predictive performance of structure C, utilizing dimensionality-reduced feature vectors in contrastive learning, outperforms structure A in AUC, F1, and MCC metrics. The superior performance of structure C can be attributed to several factors. First, the computational efficiency improves as dimension-reduced feature vectors require fewer computational resources and expedite model training. Second, it helps mitigate the risk of overfitting—a common problem in high-dimensional data—by reducing the model’s complexity. Lastly, dimension reduction aids in extracting essential features by eliminating redundant or irrelevant ones, thereby focusing the learning on the most informative aspects of the data. Therefore, despite the potential information loss, the appropriate use of dimension reduction can significantly enhance the model’s efficiency and effectiveness.

With increasing dimensionality of the feature vector, the complexity of the feature space grows, leading to elevated difficulty in contrastive learning and potentially resulting in reduced performance. As the dimensionality of the feature vector increases, redundant or irrelevant information may be introduced, which could degrade performance. Redundant or irrelevant information in high-dimensional vectors can dilute focus and introduce noise, potentially hampering model performance by obscuring key patterns. Reducing the dimensionality of feature vectors in contrastive learning can improve results for several reasons. Dimensionality reduction can help filter out these less important pieces of information, reducing noise in the model and enhancing performance. Additionally, dimensionality reduction can decrease the number of features, reducing computational complexity and making model training and inference more efficient. Furthermore, dimensionality reduction can help extract the main features of the data, preserving key information while discarding unimportant details.

However, in comparing structures B and A, we found that structure B does not necessarily outperform structure A, even though structure B utilizes dimension-reduced feature vectors during the residue-level contrastive learning stage. Upon further analysis of the model structures, we observed that the output of the model’s pooler layer needs to be dimensionally reduced by a feed-forward neural network (FNN) before it can be added to the feature vectors participating in residue-level contrastive learning. However, the loss function employed by this part of the FNN network is the cross-entropy loss function of the classification module. This may disrupt the feature vectors obtained from the type-level contrastive learning, thus introducing instability into the model. Following our comparative analysis of the three structures, structure C, which applies dimensionality reduction to feature vectors utilized in both residue-level and type-level contrastive learning, proves to be the most effective option. This superiority arises from the advantages yielded by integrating dimensionality reduction directly within the representation learning module, which significantly bolsters the model’s predictive performance and computational efficiency. Moreover, separating the representation learning module from the classification module serves as another crucial aspect of our approach. This separation ensures that the parameters of each module are updated independently, thereby preserving the integrity and stability of the learning process for each. Integrating the representation learning network into the classification module can potentially lead to instability, as the cross-entropy loss function used in the latter could distort the feature vectors derived from the type-level contrastive learning. Consequently, the independent operation of these modules fosters the preservation of significant patterns across different layers of learning, enhancing the overall stability and accuracy of our model.

### 3.3 Case Study

To further validate the performance of our model MucLiPred in predicting both DNA– and RNA-binding residues, we selected a new dataset for evaluating purposes. The selected dataset is derived from the research conducted on DRNApred^24^, a sequence-based method specifically designed for predicting DNA– and RNA-binding residues. To facilitate our discussion, we have assigned this dataset as **Dataset 3**. We have selected several protein sequences from the testing set of Dataset 3 to serve as testing cases for evaluating our model’s performance. It is important to note that during the model training phase, we still utilized the previous Merged Dataset.

Our prediction results are displayed in Figures 5A and **5B**. For the same protein sequence, we predicted its DNA binding residues and RNA binding residues. We compared our model’s predictions with the actual labels in the testing set and visualized them using PyMOL. As evidenced in the figures, our model’s predictions achieved a high degree of accuracy, closely resembling the actual labels. This suggests that during training with the Merged Dataset, the model was able to utilize all the available data effectively. When predicting DNA binding residues, it could also leverage data related to RNA binding residues that were encountered during the previous learning process, and vice versa. Moreover, during the prediction process, the model paid attention to the ligand type information, providing differing results for the distribution prediction of DNA and RNA binding residues. It’s notable that for DNA and RNA binding regions, there are some common segments. This may be related to the structure of the protein; the binding pockets of proteins are more prone to bind with ligands of the same category. Since both DNA and RNA are nucleic acids, they share some common binding regions on the protein. By introducing ligand type information, our model can distinguish between different ligand types, consciously differentiating the binding regions of different ligands. This enables better utilization of relevant information in the training dataset for more accurate predictions.

**Figure. 5.**
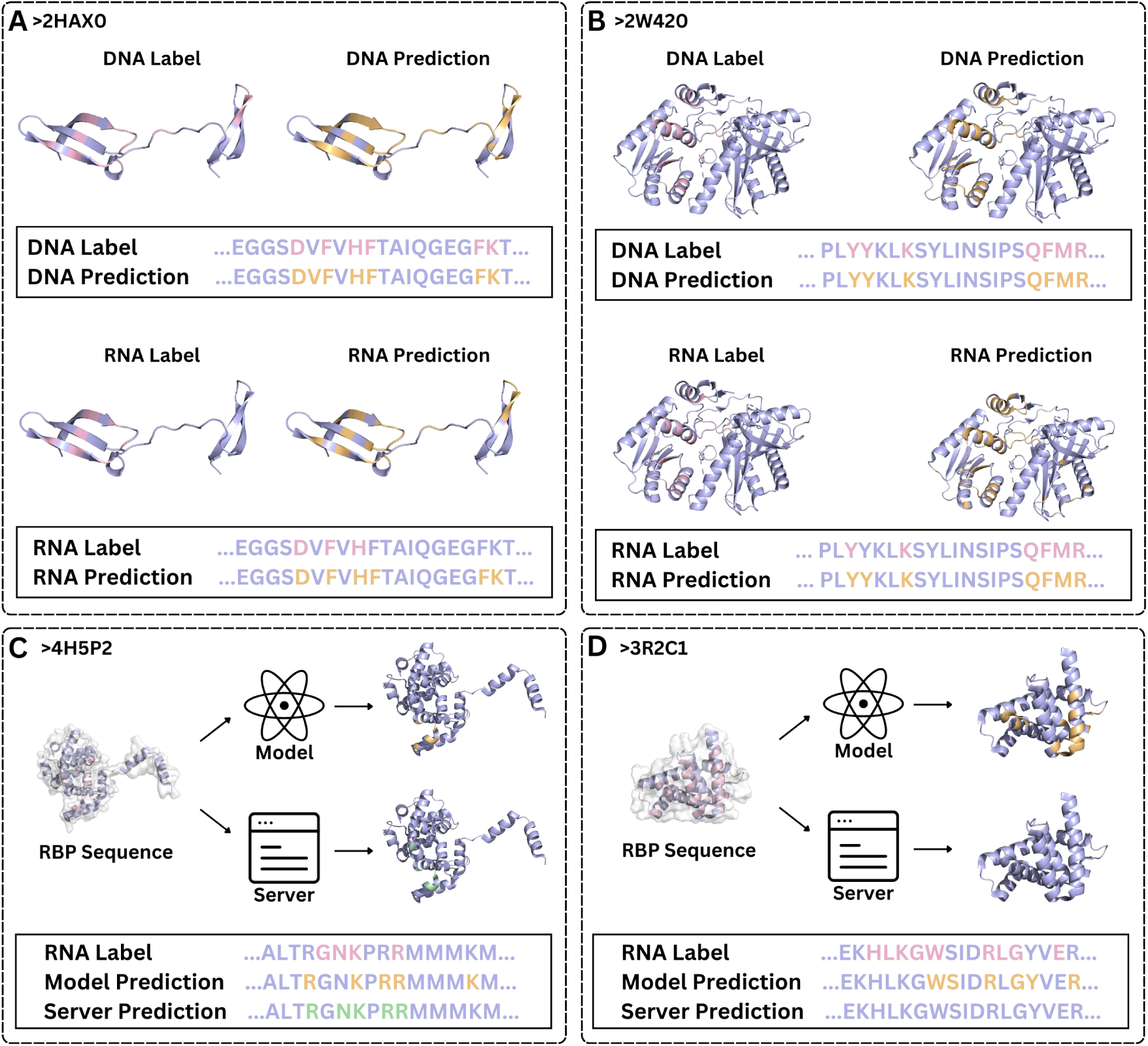
Visualization of predicted binding residues for MucLiPred. (**A-B**) Comparison between MucLiPred’s predicted DNA and RNA binding residues and the true labels from Dataset 3. **(C-D)** Comparative predictions for DNA binding residues between MucLiPred and the DRNApred web server in the absence of labels. **(C)** shows the similarity in predictions between MucLiPred and DRNApred. **(D)** illustrates a scenario where DRNApred did not predict any DNA binding residues, while MucLiPred identified some DNA binding regions.

For a specific protein sequence, it might lack the true labels for certain ligand binding residues. For such protein sequences, our model can utilize the past experience and ligand type information to provide predictions. However, evaluating the model’s predictions for these residues is challenging due to the absence of true labels. Currently, there are several publicly available webservers that can provide prediction results from their maintained models, such as DRNAPred, DeepDISOBind, and others. We used the webserver provided by DRNApred at http://biomine.cs.vcu.edu/servers/DRNApred/ as a reference. To further investigate, we modified the ligand type from RNA to DNA in the testing set of **Dataset 2** and used it as input for our model to make predictions. Concurrently, we entered the same protein sequence into the DRNApred webserver for prediction, and compared our predictions with the results from DRNApred, as shown in Figures 5C and **5D**. In Figure 5C, our predictions are similar to those of DRNApred, which serves as an indication of the validity of our model’s predictions. However, in Figure 5D, no DNA binding residues were predicted by DRNApred, while our model still identified some DNA binding regions. We believe this underscores the superiority of our model. By incorporating type-level contrastive learning, our model is capable of making more discerning predictions.

Instead of utilizing different networks to predict different types, our model operates with the same set of parameters and leverages contrastive learning to discern ligand type information and yield diverse predictions. This approach not only enhances the flexibility of our model but also reduces the computational cost and complexity, as a single network can handle different tasks. This demonstrates the power and potential of integrating contrastive learning into different protein-ligand interaction predictions, opening the door for further advancements in this field.

### 3.4 Further exploration of type-level contrastive learning

To further explore the potential of our model MucLiPred, we introduced a new ligand type – peptide. Protein-peptide interaction stands as a crucial protein interaction, holding significant importance in numerous cellular processes. Although DNA and RNA have some structural differences, they are both nucleic acids and share certain similarities. By introducing peptide into our study, with its structure being short chains of amino acids linked by peptide bonds, we can investigate the performance of our model on ligands with significantly different molecular structures.

The dataset of peptide-binding proteins was initially introduced in the research of the structure-based method SPRINT-Str^37^ and is referred to as **Dataset 4** in this paper. As with the previous operation for generating the Merged Dataset, we separately mixed the training sets of Dataset 1 and Dataset 2 with the training set of Dataset 4, removing some data to ensure an equal number of samples for the two ligand types. The same preprocessing was applied to their respective testing sets as well. The dataset resulting from the combination of Dataset 1 and Dataset 4 is referred to as **DP Dataset**, while the dataset obtained by combining Dataset 2 and Dataset 4 is referred to as **RP Dataset**. In our model, the loss function comprises three components: L_1, representing residue-level contrastive learning loss, L_2, representing type-level contrastive learning loss, and L_3, representing the cross-entropy loss in the classification module. By controlling the weights of each loss, we assign them different degrees of importance. The type-level contrastive learning loss, L_2, plays a crucial role in our model’s ability to distinguish between different ligand types.

Nowadays, many research works have been proposed for interpreting the mechanisms behind the deep learning models in the field of bioinformatics ^38, 39^. To delve deeper into our introduced type-level contrastive learning, we employed UMAP (Uniform Manifold Approximation and Projection) to visualize the output of our model after dimensional reduction. UMAP, a powerful dimensionality reduction technique, preserves the global structure of data, making it an excellent tool for visualizing high-dimensional data in two or three dimensions. By mapping the high-dimensional output of our model into a 2D space, we can visually evaluate how well our model is distinguishing between different ligand types, as shown in Figures 6A and **6B**. We have termed the weight of the type-level contrastive learning loss L_2 as the type loss weight (or w_2). When the type loss weight is zero, data points overlap and become indistinguishable, signifying that the model fails to significantly differentiate these categories in high-dimensional space. Interestingly, once we escalate the type loss weight to 0.1, the differently colored points in the UMAP plot segregate clearly, indicating that the model is efficiently differentiating the ligand types in high-dimensional space and taking into account the ligand type information. Further increasing the type loss weight to 0.5 results in an even clearer distinction between the differently colored points in the UMAP plot, and points representing the same ligand type move closer together. Simultaneously, as depicted in Figures 6C and **6D**, there is a consistent rise in the three indicators – Precision, F1 score, and MCC – with the progressive increase of the type loss weight. The upliftment of the type loss weight from 0.0 to 0.1 results in a notable enhancement in the model’s performance. A minute alteration can bring about a substantial improvement, demonstrating the rationality of our model design and its immense potential.

**Figure. 6.**
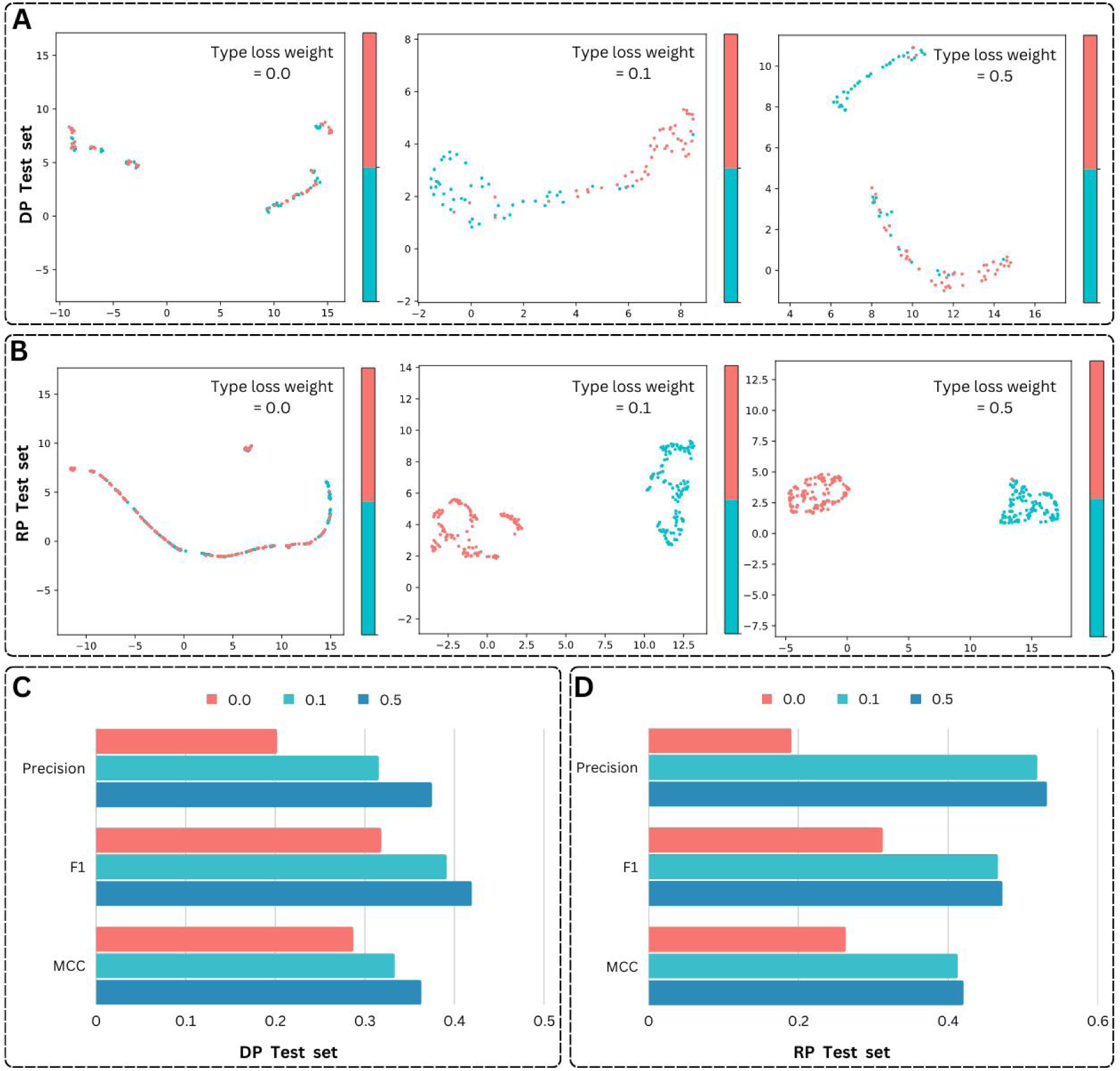
Illustration of type-level contrastive learning effects. (**A-B**) UMAP plots illustrate the feature distribution of proteins binding with different ligand types (marked by different colors) in 2D space, for different datasets, with increasing type loss weights (w_2). **(C-D)** Evaluation of Precision, F1 score, and MCC for different datasets, with increasing type loss weight (w_2). The corresponding raw data is also available in **Supplementary Table 3**.

The model uses the training set comprising different ligand-protein binding information for two distinct ligand types. By incorporating type information into the input, our model is capable of simultaneously learning the distribution of binding residues for these two different ligand types. The learned representations push apart different categories, while pulling closer the same categories, allowing for accurate prediction of the binding sites for each ligand type. This ability to learn and focus on the differences between various ligand types by adjusting the type loss weight demonstrates the power and adaptability of our model. It highlights the importance and benefits of incorporating type-level contrastive learning in predicting protein-ligand binding residues, showcasing the potential for enhancing the performance in various protein-ligand interaction prediction tasks. Expanding our model to include ligand categories with considerably different molecular structures further demonstrates the adaptability and robustness of our architectural design. The ability to accurately predict the binding residues across such diverse ligand categories underscores the model’s generalizability and its potential to be applied broadly across various protein-ligand interaction prediction tasks.

Moreover, the integration of type-level contrastive learning is particularly advantageous when dealing with ligands that are structurally diverse. It allows the model to capture distinct, type-specific patterns and characteristics, thereby improving the quality of the learned representations and enhancing prediction performance. In essence, our model’s capacity to handle and learn from such structurally diverse ligand types underscores its versatility, promising its utility in future protein-ligand interaction studies of diverse nature.

## 4 Discussion

Protein-ligand interactions play a critical role in many biological functions, making their accurate prediction pivotal for drug discovery and design processes. In our research, we have developed a dual contrastive learning approach, named MucLiPred, which is utilized to simultaneously predict different protein-ligand binding residues exclusively using protein sequences. The benchmarking experiments provide evidence that the proposed MucLiPred outperforms existing single-type and multi-type prediction methods across all metrics.

Specifically, our approach employs the widely adopted pre-training model BERT as the encoder, facilitating automatic learning and generation of enhanced representations for protein sequences. We fully exploit the output of the Pooler layer as the feature vector for type-level contrastive learning. The use of Pooler output captures the global semantic representation of the entire sequence, which is crucial for tasks that require comprehensive sequence-level information.

For our classification module, we employ a process known as ‘addition’ to fuse information. In the realm of deep learning, information fusion refers to the integration of information from diverse sources or hierarchical levels. We incorporate ligand type information of the sequence into the output of the Encoder within the classification module. The superior performance of the addition-based fusion in our case could be attributed to its ability to blend feature spaces and establish direct interactions between features. In a further innovative step, our model incorporates a type-level contrastive learning mechanism. The contrastive learning paradigm operates at the type level, pushing representations of different ligand types apart and pulling similar ones closer. This approach has proven effective, as evidenced by our use of UMAP for visual evaluations. The adjustment of the weight of type-level contrastive loss facilitates the model’s adaptability, resulting in significant improvements in metrics like Precision, F1 score, and MCC.

However, while our current strategy has been effective, there are still opportunities to further optimize components and enhance performance. A future extension could be to predict the strength of the binding interaction, providing an even more nuanced understanding of protein-ligand interactions. Looking forward, the capacity of our proposed MucLiPred model in predicting protein-ligand binding residues represents a significant advancement in this field. As we further refine and extend this approach, it promises to have a substantial impact on fields like bioinformatics, drug discovery, and molecular biology.

## Availability of Data and Materials

The data and code for this study can be found in a GitHub repository accompanying this manuscript:

## Author contributions

J.Z. conceived the basic idea and designed the framework. J.Z. and R.W. performed the experiments. J.Z. and R.W. wrote the manuscript. L.W. revised the manuscript. J.Z. conducted the visualization of the experimental results.

