## Supplementary Materials for "Multi-Level Contrastive Learning for Protein-Ligand Binding Residue Prediction"

### Supplementary Tables

**Supplementary Table 1. The experimental results under different architectures on the testing set of Dataset 1**

|  | Feature vectors | | | Fusion approach | | | Structure | | |
| --- | --- | --- | --- | --- | --- | --- | --- | --- | --- |
|  | Mean | None | Pooler | Concat | None | Add | A | B | C |
| AUC | 0.836 | 0.855 | 0.859 | 0.854 | 0.849 | 0.859 | 0.852 | 0.854 | 0.859 |
| F1 | 0.398 | 0.302 | 0.485 | 0.376 | 0.467 | 0.485 | 0.463 | 0.369 | 0.485 |
| MCC | 0.364 | 0.285 | 0.436 | 0.345 | 0.413 | 0.436 | 0.414 | 0.344 | 0.436 |

**Supplementary Table 2. The experimental results under different architectures on the testing set of Dataset 2**

|  | Feature vectors | | | Fusion approach | | | Structure | | |
| --- | --- | --- | --- | --- | --- | --- | --- | --- | --- |
|  | Mean | None | Pooler | Concat | None | Add | A | B | C |
| AUC | 0.753 | 0.833 | 0.843 | 0.801 | 0.816 | 0.843 | 0.802 | 0.811 | 0.843 |
| F1 | 0.317 | 0.462 | 0.513 | 0.464 | 0.481 | 0.513 | 0.485 | 0.489 | 0.513 |
| MCC | 0.202 | 0.381 | 0.430 | 0.368 | 0.383 | 0.430 | 0.399 | 0.393 | 0.430 |

### Supplementary Table 3. **The experimental results** for DP and RP datasets, with increasing type loss weight.

|  | DP test set | | | RP test set | | |
| --- | --- | --- | --- | --- | --- | --- |
|  | 0.0 | 0.1 | 0.5 | 0.0 | 0.1 | 0.5 |
| Precision | 0.202 | 0.315 | 0.375 | 0.191 | 0.519 | 0.532 |
| F1 | 0.318 | 0.391 | 0.419 | 0.312 | 0.466 | 0.472 |
| MCC | 0.287 | 0.333 | 0.363 | 0.263 | 0.412 | 0.420 |
